# Energy-precision trade-off in mitotic oscillators revealed by ATP modulation in artificial cells

**DOI:** 10.64898/2026.03.02.709190

**Authors:** Shiyuan Wang, Liam Yourston, Gembu Maryu, Yeonghoon Kim, Derek Walker, Usha Kadiyala, Qiong Yang

**Author notes:** These authors contributed equally (Shiyuan Wang, Liam Yourston). These authors contributed equally (Gembu Maryu, Yeonghoon Kim).

## Abstract

The temporal precision of biochemical oscillators is fundamentally constrained by the energy dissipated to suppress molecular fluctuations, a widely predicted trade-off governing information processing across biology and physics, from molecular motors to kinetic proofreading to computing. Yet, experimental validation in complex biological oscillators remains elusive due to challenges of systematically modulating energy while quantifying stochastic dynamics across large ensembles. Here, we establish a high-throughput droplet-microfluidics platform to reconstitute mitotic oscillations from *Xenopus laevis* egg extracts within thousands of sub-nanoliter compartments. By precisely tuning ATP across a broad free-energy landscape and developing an analytical framework that decouples intrinsic phase diffusion from quenched period heterogeneity, we uncover a hidden trade-off linking metabolic budget, oscillation speed, and precision. While speed peaks non-monotonically near physiological ATP levels and declines toward both high and low bifurcation limits, precision increases monotonically with energy. These findings provide direct experimental evidence that mitotic timing is actively shaped by energy budgets. Intriguingly, embryonic cell cycles are not optimized for maximum fidelity, but for a metabolic compromise maintaining just enough coherence for synchronous yet rapid divisions, placing the endogenous ATP budget near an energetic optimum balancing speed and accuracy. Our integrated artificial-cell and analytical strategy provides a generalizable framework for mapping thermodynamic limits in non-equilibrium biological dynamics.

---

Early embryos of many species, including *Xenopus* and zebrafish, undergo exceptionally rapid and synchronous cleavage cycles that run without transcription or cell growth and rely solely on maternally deposited resources. Within hours after fertilization, a single egg partitions its cytoplasm into hundreds to thousands of progressively smaller cells with striking temporal accuracy, approximately every 30 minutes in *Xenopus laevis* and every 15 minutes in zebrafish, with a period coefficient of variation (CV) below 5%^1–3^. This combination of high precision, high speed, and strict energetic limitation highlights the robustness and efficiency of the underlying oscillator and raises a fundamental question: how do embryonic cell-cycle clocks maintain accurate timing under metabolic constraints?

These divisions are driven by a cyclin-dependent kinase 1 (Cdk1) oscillator conserved across eukaryotes (Fig. 1**a**, top). Cyclin B synthesized from maternal mRNA binds to Cdk1, whose activation is regulated by dual positive feedback through the opposing Cdc25 phosphatase and Wee1 kinase, creating a bistable switch at mitotic entry^4–6^. Active Cdk1 then triggers cyclin B degradation via the anaphase-promoting complex/cyclosome (APC/C), completing a negative feedback loop that resets the cycle^7,8^.

**Fig. 1.**
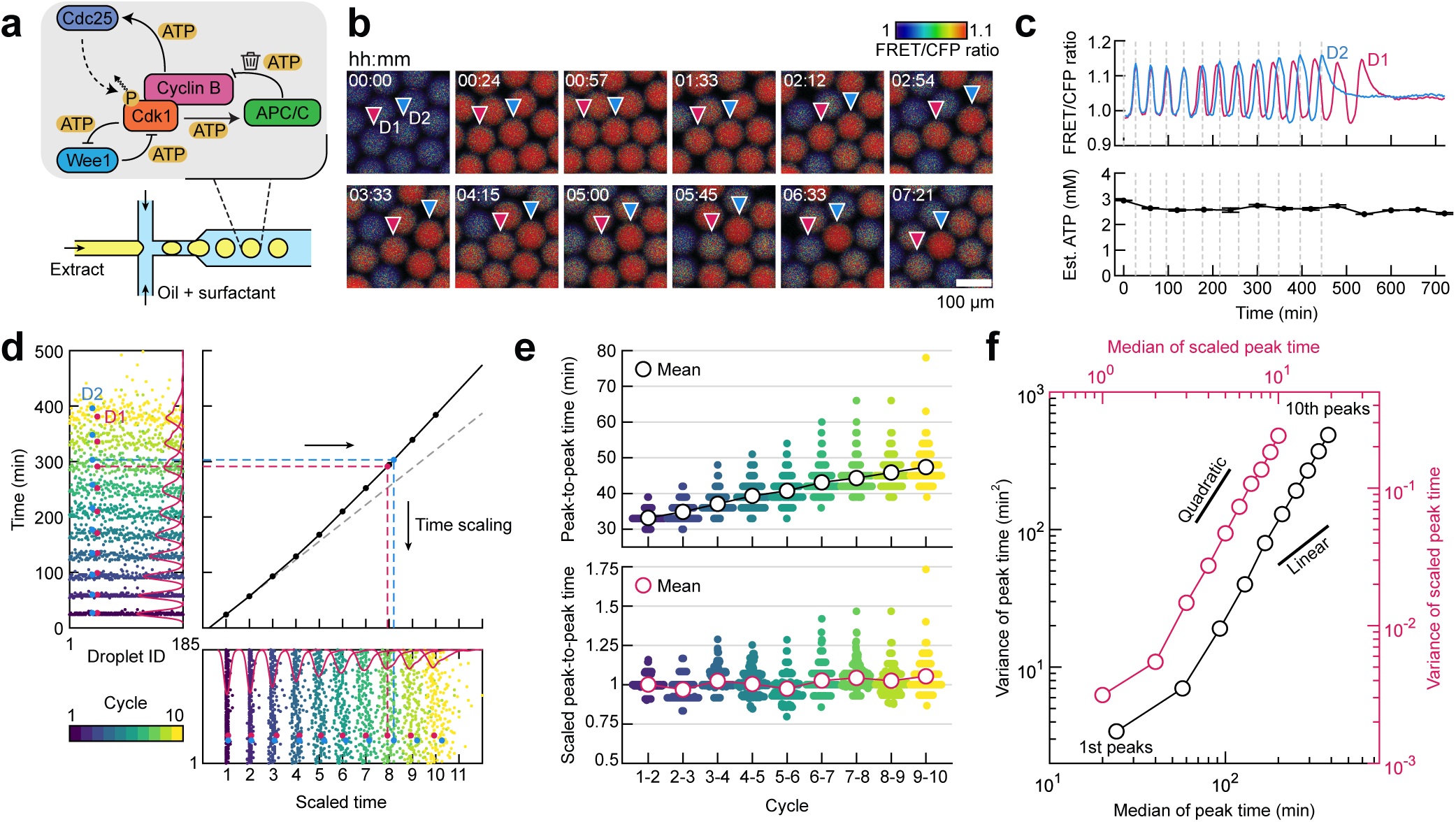
Phase diffusion in compartmentalized cell-free extract. **a** Top: Schematic of the Cdk1 oscillator network, formed by coupled positive and negative feedback loops, with ATP-dependent phosphorylation–dephosphorylation reactions fueling each regulatory step. Bottom: Single-aqueous-input flow-focusing microfluidic device for encapsulating extracts that contain the Cdk1 network into monodisperse water-in-oil microemulsion droplets. **b** Time-lapse montage of a representative field of view showing droplets at the indicated times (hh:mm). The FRET/CFP emission ratio is rendered in intensity-modulated display mode, with color indicating Cdk1 activity. The color bar ranges from low (blue) to high (red). Two neighboring droplets, D1 (red triangle) and D2 (blue triangle), are randomly selected to illustrate dephasing over time. Scale bar: 100 µm. **c** Top: Time courses of spatially averaged FRET/CFP ratio for droplets D1 and D2 indicated in **b**. Bottom: Time course of estimated endogenous ATP concentration measured concurrently by a luciferase assay. Vertical dashed lines mark corresponding time points in the montage in **b**. Error bars: standard deviation from three technical replicates. **d** Left: Raster plot of oscillation peak times across the first 10 cycles for an ensemble of *N* = 185 independent droplets. Droplets D1 and D2 from **b**-**c** are highlighted in their respective colors. Solid red lines trace the distribution of the peak times for each cycle. Middle: Time scaling. A reference scaling function (solid black line) is defined as a piecewise linear interpolation of {(*T* ^0^, *n*)} (black dots), where *T* ^0^ is the median *n*-th peak time. The red and blue dashed lines represent the scaling trajectories for D1 and D2, respectively. The dashed gray line is the linear extrapolation from the first two cycles. Bottom: Raster plot of scaled peak times. **e** Top: Distribution of time intervals between successive peaks, demonstrating overall period lengthening over cycles. Bottom: Distribution of peak-to-peak time intervals after scaling. **f** Variance of raw (scaled) peak times as a function of raw (scaled) peak time median, illustrated in black (red). The variance increases quadratically over time regardless of scaling.

Sustaining this oscillator far from equilibrium demands continuous ATP hydrolysis at each regulatory step, from multisite phosphorylation of Cdk1, Wee1, Cdc25, and APC/C^9,10^ to cyclin B synthesis and ubiquitin-proteasomal degradation^11–13^. Beyond the intrinsic cost of the clock itself, Cdk1 activation triggers a cascade of downstream processes that impose a substantial extra energetic “load”. Cdk1 phosphorylates hundreds of substrates (*>* 1, 000 sites)^14–16^ and orchestrates numerous ATP-intensive mitotic events, including DNA replication^17^, chromosome condensation^18^, nuclear envelope breakdown and reformation^19,20^, spindle assembly^21^, centrosome positioning^22^, and sister chromatid separation^23^. Consistently, calorimetric measurements in *Xenopus* and zebrafish embryos reveal heat oscillations synchronized with cleavage cycles and peaking at mitotic entry, confirming mitosis as the most energy-intensive phase^24,25^. With finite maternal ATP distributed across dozens of rapid cycles, these observations imply tight coupling between mitotic dynamics and its energetic environment.

Theoretical work by Cao *et al.* suggested that excess free-energy dissipation beyond an oscillator’s basal requirements suppresses phase fluctuations by enforcing net forward flux through state space, imposing a thermodynamic cost of precise periodicity^26^. Specifically, the phase diffusion constant *D*, which describes the growth rate of peak-time variance, is predicted to decrease with increasing dissipation per cycle, exemplifying a biochemical instance of the thermodynamic uncertainty relation^27^. Intriguingly, ATP levels in cleaving embryos are significantly higher than in adult tissues and decline as development progresses^28^, suggesting that elevated maternal ATP may be a physical requirement for sustaining early embryonic cycle precision and synchrony before zygotic metabolic controls emerge^29^. Despite its profound implications, this predicted energy-precision trade-off has lacked direct experimental validation, due to the difficulty of disentangling intrinsic energy cost from downstream “loads” and of systematically controlling ATP availability over a wide dynamic range *in vivo*.

Here, we overcome these limitations by reconstituting a minimal mitotic oscillator in ATP-controlled, high-throughput microfluidic droplets. These sub-nanoliter artificial cells contain the complete Cdk1 oscillator network from the *Xenopus laevis* egg cytoplasm but exclude nuclei and other major ATP-intensive downstream processes (e.g., DNA replication). The sub-nanoliter droplet volume (*V*_droplet_) ensures rapid diffusive equilibration 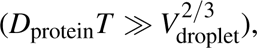 allowing each droplet to be treated as a homogeneous, zero-dimensional reactor with a well-defined oscillator period (*T*). We formulate a generalized analytical framework to extract the intrinsic phase diffusion constant *D* from ensemble dephasing, yielding dimensionless *D/T* on the order of 10^−3^–10^−2^, a precision sufficient to maintain the observed synchrony of the first 12 cleavages in *Xenopus* embryos before the mid-blastula transition (MBT). Moreover, ATP availability negatively correlates with mitotic timing noise, as demonstrated by decreasing *D* upon exogenous ATP addition and increasing *D* upon enzymatic depletion. Taken together, our findings provide direct experimental evidence that cellular free-energy budget serves not merely as a fuel but as an active regulator of mitotic timing, and that early embryos operate under thermodynamic constraints that enforce an optimal balance between energetic efficiency, speed, and temporal precision.

## Mitotic oscillators exhibit superlinear phase diffusion in droplet ensembles

To quantify the precision of mitotic oscillations under controlled energetic conditions, we utilized flow-focusing microfluidics to encapsulate cycling *Xenopus laevis* egg extracts^30^ within monodisperse, cell-like droplets (∼ 100 µm in diameter; Fig. 1**a**, bottom). Each droplet functions as an independent biochemical oscillator, containing the core machinery of Cdk1 network (Fig. 1**a**, top), a Cdk1 FRET biosensor (Cdk1-EV^31^), and where indicated, an exogenous creatine/phosphocreatine energy regeneration system (energy mix). Real-time Cdk1 dynamics were tracked as the mean FRET/CFP ratio within each droplet (Fig. 1**b**; Fig. 1**c**, top), at a temporal resolution of 3–6 min. Cdk1 oscillations were initially synchronized across droplets prepared from a homogenized bulk extract, but gradually lost coherence over successive cycles (Fig. 1**b**; Fig. 1**c**, top; Supplementary Video 1). Raster plots of a population of droplets (*N* = 185) (Fig. 1**d**, left) revealed progressive ensemble dephasing of *n*-th peak times, evident as a gradual broadening of their distribution (Fig. 1**d**, left, red lines).

This loss of synchrony was accompanied by systematic period lengthening (Fig. 1**e**, top). We previously hypothesized that ATP depletion drives such slowing in cell-free systems^32^. However, time-course luciferase assays revealed stable ATP concentrations maintained at endogenous levels (∼ 3 mM, within reported physiological ranges for amphibian oocytes and eggs^33,34^) throughout and beyond the oscillator lifetime (Fig. 1**c**, bottom), or higher with energy mix supplementation (Fig. S1). This finding confirms previous reports^35^ that cytoplasmic extracts preserve robust ATP homeostasis through metabolism of endogenous glycogen stores, thereby providing a foundation for precise control of the energy budget in droplet-based artificial cells. In contrast, RT-qPCR revealed degradation of maternally deposited cyclin B1 mRNA that correlated with cell cycle elongation and termination (Fig. S2). Together with our previous findings that cyclin B1 mRNA levels directly tune cell cycle periods^31,36^, we conclude that the observed slowdown is primarily due to reduced cyclin B synthesis kinetics, not energy depletion.

Such non-stationary behavior poses the first challenge for phase fluctuation analysis, as classical models are formulated exclusively for stationary dynamics^26^. While these models predict linear growth of peak-time variance (*σ* ^2^ ∝ *t*)^26^, our ensemble data showed superdiffusive scaling, with variance increasing approximately quadratically over time (Fig. 1**f**, black). To determine whether this anomaly was an artifact of period elongation, we rescaled time to equalize the intervals between consecutive median peak times (Fig. 1**d**, bottom; Supplementary Information Section 1). Remarkably, even after this normalization (Fig. 1**e**, bottom), both the gradual dephasing (Fig. 1**d**, bottom) and the quadratic variance growth (Fig. 1**f**, red) persisted, indicating that dephasing is governed by more than simple Brownian phase noise, necessitating a generalized analytical framework to accurately resolve the oscillator’s intrinsic precision.

## A stochastic framework isolates intrinsic phase diffusion from ensemble heterogeneity

Existing precision-dissipation theory has mainly focused on intrinsic stochasticity, an inherent randomness of molecular collisions and reactions that becomes significant when molecular species are in low copy numbers^26,37^. However, ensemble dephasing can arise from both intrinsic stochasticity of statistically identical oscillators [Fig. 2**a** (i)] and extrinsic factors, such as heterogeneity in oscillator periods [Fig. 2**a** (ii)]. The stochastic partitioning of bulk reactions into sub-nanoliter compartments, analogous to the “partitioning errors” during natural cell divisions^38^, can introduce “quenched randomness” in molecular composition, leading to diverse dynamical phenotypes, such as period variability that often exhibit super-Poissonian statistics^39^. Critically, such heterogeneity in compartmentalized systems emerges independently of energy dissipation by the underlying molecular circuitry, yet can dominate ensemble dephasing and overshadow the dissipation-dependent intrinsic noise governed by metabolic constraints [Fig. 2**a** (iii)]. Therefore, we developed a framework to disentangle intrinsic stochasticity from period heterogeneity, enabling accurate analysis of energetic constraints on oscillator precision.

**Fig. 2.**
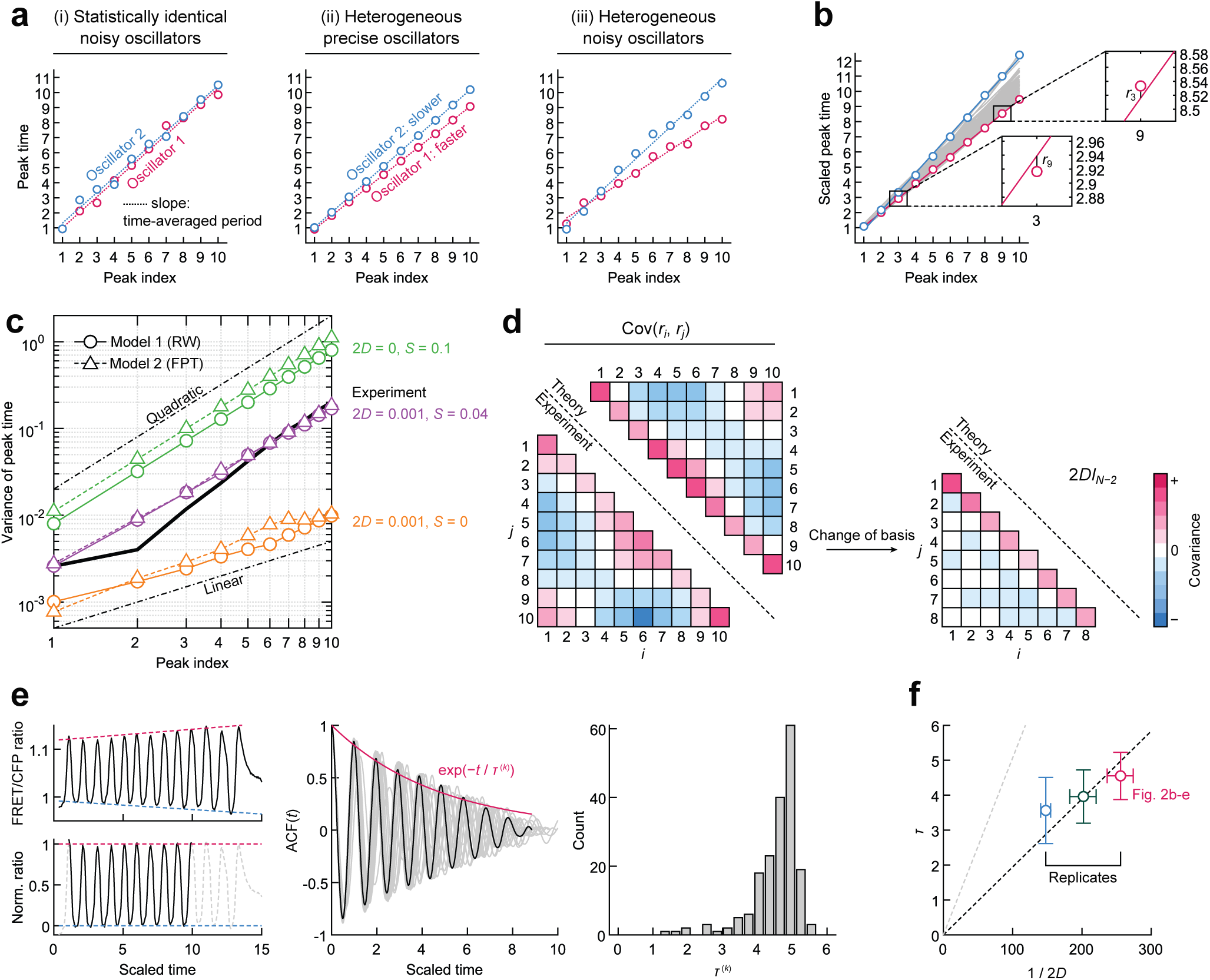
Quantification of phase fluctuations. **a** Ensemble dephasing can arise from two sources: (i) intrinsic noise and (ii) heterogeneous oscillator dynamics. (iii) exemplifies a situation where heterogeneity in periods (slopes of linear trend lines) dominates intrinsic stochasticity. **b** Peak time versus peak index for the mitotic oscillators presented in Fig. 1 (gray lines). The oscillator with the shortest (longest) time-averaged period is indicated in red (blue). Insets exemplify the residual peak time noise relative to the time-averaged projection. **c** Nonlinear scaling of peak time variance reproduced with two stochastic models. Examples are provided for three (*D, S*) conditions (in orange, green, and purple colors) for each model (markers). The experimental scaling relationship (Fig. 1f) is replotted for comparison (solid black line). **d** *D* estimation. Left: Covariance matrix of residuals for the first *N* = 10 peak times (lower triangle, insets of Fig. 2b), compared with theoretical reference (upper triangle, Supplementary Information Section 2). Right: After standardization, the covariance matrix (lower triangle) is expected to be an identity matrix multiplied by 2*D* (Supplementary Information Section 2). **e** Autocorrelation function (ACF) analysis. Left: The amplitude of a raw FRET/CFP ratio time series (top) is normalized (bottom) by scaling its peak and trough intensity trends (dashed red and blue lines). To ensure the consistency with *D* estimation, a truncated time series between the first and *N*-th peaks (bottom, solid black line) is used for computing the ACF. Middle: ACFs of the illustrated example (black line) and 18 other randomly selected time series (gray lines). The exponential decay of the autocorrelation function defines the characteristic dephasing time *τ*^(^*^k^*^)^ for a single droplet oscillator. *τ*^(^*^k^*^)^ = 4.72 for the example. Right: *τ*^(^*^k^*^)^ histogram. The ensemble-level correlation time *τ* = 4.55 is defined as the mean of *τ*^(^*^k^*^)^. **f** Reciprocal relationship between *D* and *τ* for three biological replicates. An estimated coefficient is 0.02 (dashed black line), whereas (2*π*^2^)^−1^ is expected for sinusoidal oscillations (dashed gray line). Horizontal error bars: 95% confidence intervals as defined in Supplementary Information Section 2. Vertical error bars: standard deviation of single-droplet *τ*^(^*^k^*^)^.

Our droplet data confirmed that period heterogeneity is significant (Fig. S3) and is the primary source of the observed population dephasing. Scaled peak times of individual droplets plotted against cycle index (Fig. 2**b**; data taken from Fig. 1) revealed near-linear scaling, suggesting that an individual oscillator can be characterized by a near-constant period with modest fluctuations in peak-to-peak intervals. Nevertheless, variations in periods across the droplet population led to progressive broadening of the ensemble peak-time distribution (Fig. 1**d**). To reproduce this behavior, we developed two asymptotically equivalent stochastic models (Supplementary Information Section 2), characterized by two parameters: the intrinsic diffusion constant *D* and the period standard deviation *S*, both nondimensionalized by time rescaling (Fig. S4**a**). One model emulates a peak time sequence as Gaussian random walks (RW) under a constant drift (Fig. S4**b**, left), and the other generates peak times as first-passage times (FPT) of a drifted Brownian particle to reach successive thresholds set at equidistant intervals (Fig. S4**b**, right). In the limit of negligible period heterogeneity (2*D* ≫ *S*^2^), peak-time variance increased linearly over time with slope 2*D* (Fig. 2**c**, orange). In contrast, significant period heterogeneity (2*D* ≪ *S*^2^) resulted in nonlinear variance growth (Fig. 2**c**, green; Fig. S4**c**). Consistent with experimental observations (Fig. 2**b**), the data (Fig. 2**c**; solid black line) were best captured by the models in the heterogeneity-dominated regime (Fig. 2**c**, purple, 2*D < S*^2^).

We extracted *D* from the residual noise of peak-to-peak intervals relative to the time-averaged period (Fig. 2**b**, inset). RW model analytically shows that the covariance of the residual noise depends solely on *D* and is insensitive to *S* (Supplementary Information Section 2). Thus, *D* can be estimated by comparing the experimental covariance matrix to theoretical predictions (Fig. 2**d**), irrespective of the heterogeneity level *S*. Validation using synthetic time series data from both stochastic models demonstrated the robustness and accuracy of this generalized estimator (relative error *<* 4.4%; Fig. S4**d**). Application of the estimator to the experimental data yielded 2*D* = 3.9 × 10^−3^, comparable in order of magnitude to the theoretical lower bound predicted for a simple activator-inhibitor model^26^.

As an independent alternative validation of the estimator, we measured precision using the correlation time of Cdk1 oscillations (Supplementary Information Section 3). The correlation time *τ* was obtained as the population mean of the characteristic decay time *τ*^(^*^k^*^)^ of the envelope of the normalized FRET/CFP ratio autocorrelation for individual droplets (Fig. 2**e**). The independent quantification of *D* and *τ* across three biological replicates exhibited the anticipated inverse relationship, with the proportionality constant 0.02 (Fig. 2**f**). The deviation of this constant from the theoretical reference (2*π*^2^)^−1^ `≈` 0.05 can be attributed to the spiky, rather than sinusoidal, waveform of Cdk1 activity^26^.

## Systematic ATP tuning reveals non-monotonic speed response and an energy-precision trade-off in mitotic oscillator

To examine how energy availability influences oscillator performance, we first manually varied energy mix concentrations (0.5–3.5 mM) in bulk extracts to generate droplet populations at discrete ATP levels (Fig. 3**a**). While our analytical framework revealed a consistent reduction in the phase diffusion constant 2*D* with increasing free energy (Fig. 3**b**), variability across biological replicates and the coarse, discrete nature of manual titrations hindered accurate resolution of the continuous ATP dependence of oscillation dynamics (Fig. 3**c**). Finite sample preparation times further complicated identification of the true first oscillation cycle under faster conditions (Fig. 3**a**, black dots), potentially introducing misalignment of peak indices across ATP levels. These limitations necessitate a high-throughput, continuous ATP tuning strategy to map the complete dynamical response.

**Fig. 3.**
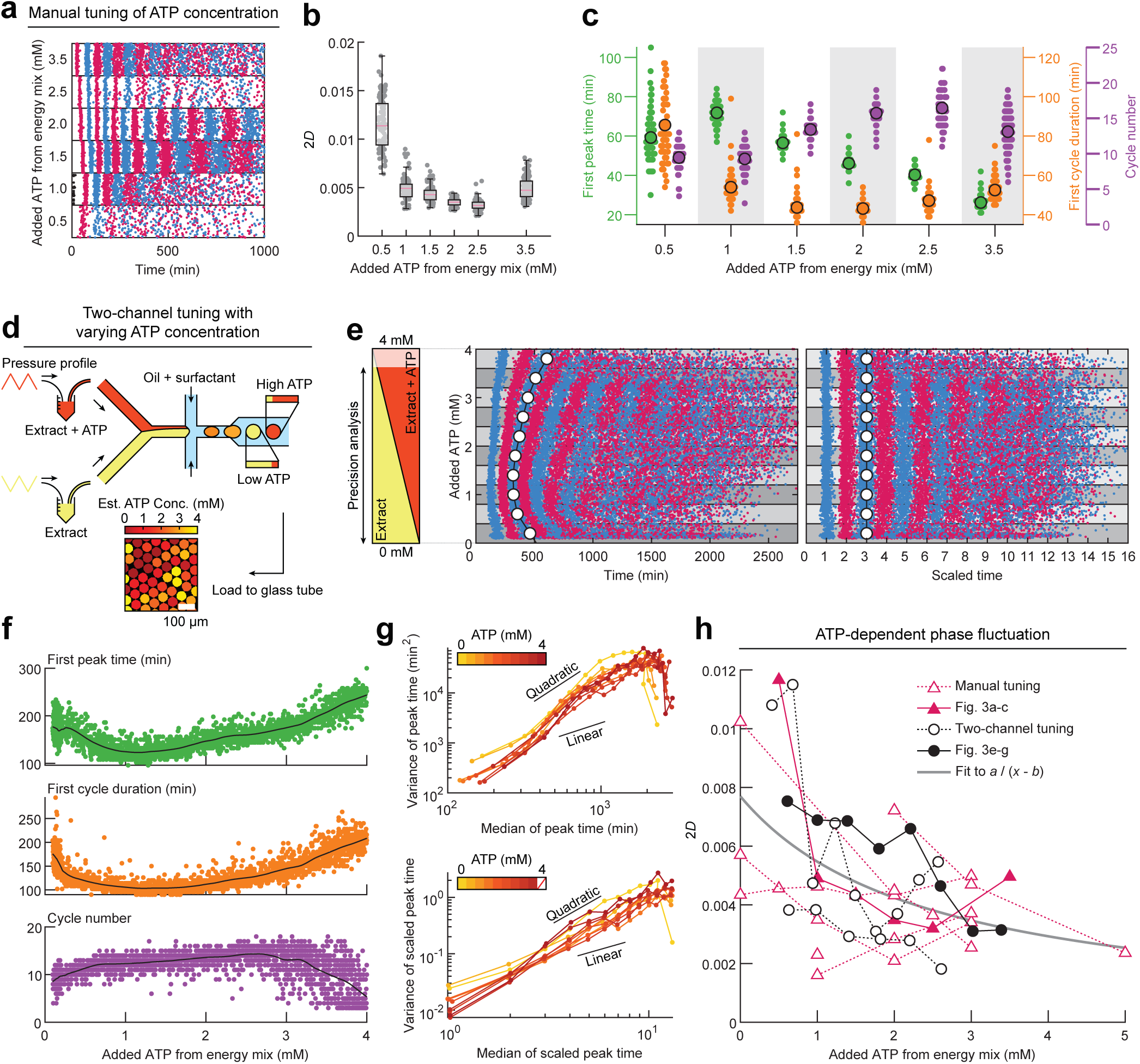
Systematic exogenous ATP tuning reveals that energy availability shapes mitotic oscillator speed and precision. **a** Representative raster plots for manually tuned ATP conditions using a single extract preparation. **b** Estimated 2*D* for the corresponding data in **a**. Stationary criteria of *R*^2^ *>* 0.95 were applied. Horizontal red lines indicate median, boxes the interquartile range, and whiskers the 0.05–0.95 quantile. **c** Corresponding quantification of first peak times (as defined in **a**), first peak-peak period, and cycle number for the data in **a**. **d** A two-aqueous-inlet flow-focusing microfluidic device is used to generate droplets by driving the extract inlets with opposite pressure waveforms to tune droplet composition (top). The ATP content of each droplet is estimated from the fluorescence intensity of an inert fluorescent dye introduced through one of the extract inlets, as shown in a representative field of view of the droplets (bottom). **e** Data is plotted based on estimated ATP concentration. Droplets are binned based on ATP levels to generate a standard curve calculated from the median peak-times (white). The peak time raster plot as a function of ATP (left) is mapped onto integer cycle numbers (right). **f** Time of the first cycle peak time as observed from two-channel data (top), plotted based on the droplet’s tracking dye intensity to represent added ATP. The period of the first cycle demonstrates a non-monotonic response in cell cycle timing due to ATP levels (middle). The number of peaks observed in each droplet (bottom). **g** Plots of the non-linear trend in peak-variance for the binned data in **e**. Peak variance is suppressed at intermediate ATP levels for real-time values (top) as well as time-rescaled data (bottom). **h** 2*D* from ATP tuning across multiple biological replicates. Stationary criteria of *R*^2^ *>* 0.95 were applied to all datasets.

To rigorously map the oscillator response across a continuous energy gradient, we developed a two-channel microfluidic tuning system for high-throughput production of oscillators across controlled conditions. Extracts supplemented with either no or high-concentration energy mix were injected through pressure-controlled inlets and mixed at varying ratios prior to encapsulation (Fig. 3**d**, top). ATP concentration in each droplet was determined by the fluorescence intensity of an inert dye co-injected at the high-ATP inlet (Fig. 3**d**, bottom). This ATP quantification was further validated by independent measurements using a ratiometric ATP biosensor QUEEN-2m^40^ (Fig. S5**a**).

Raster plots from *N* = 2, 481 droplets revealed a non-monotonic dependence on ATP (Fig. 3**e**, left; Supplementary Video 2), with an optimal intermediate ATP range enabling earlier first peak time, faster initial speed, more cycle number as compared to low and high boundaries (Fig. 3**f**). These observations suggest that the supplemented energy mix, capable of sustaining long-term ATP homeostasis in our extracts (Fig. S1), can modulate both early-cycle kinetics and long-term oscillator stability.

With periods now ATP-dependent, we performed time rescaling using standard curves of the *n*-th peak times from binned two-channel datasets (Fig. 3**e**, white), which served as the scaling reference for individual oscillator peak times. This transformation largely removed ATP-dependent period heterogeneity, yielding near-uniform periods across ATP conditions (Fig. 3**e**, right). To ensure *D* estimates reflected intrinsic stochastic phase fluctuations rather than near-bifurcation instabilities, we applied exclusion criteria based on cycle number and linearity of scaled peak times (Fig. S6; Fig. S7; Supplementary Information Section 4). This step removed anomalous time series exhibiting accelerated period lengthening in their final cycles near termination, a behavior characteristic of critical slowing near bifurcation, likely triggered by the degradation and exhaustion of maternal cyclin B mRNA (Fig. S2**c**) or proximity to ATP low or high extremes (Fig. S7**c**). Final scaled data were obtained by reapplying time rescaling exclusively to the stationary, well-behaved population.

Consistent with the untuned condition (Fig. 1**f**), peak-time variance exhibited nonlinear growth both before and after time rescaling across all ATP conditions (Fig. 3**g**), confirming that period heterogeneity dominates the apparent dephasing over intrinsic noise. Variance was minimized at intermediate ATP, reflecting the nonmonotonic period response. However, by applying our covariance-based estimator to segregate the intrinsic noise *D* from period heterogeneity, a clear thermodynamic trade-off between timing accuracy and free-energy dissipation emerged: oscillator precision increased (*D* decreased) monotonically with ATP concentration (Fig. 3**h**, black line), a consistent trend held across independent replicates and manual titration controls (Fig. 3**h**, magenta). Notably, this suppression of phase fluctuations persisted even without phosphocreatine (CP) (Fig. S8), suggesting that the initial ATP deposit, sustained by endogenous homeostatic mechanisms, is sufficient to modulate oscillator precision over extended times.

These findings provide direct experimental evidence that embryonic cell cycle operates under a thermodynamic trade-off, where cellular energy status controls the fidelity of cell-cycle timing and ATP mechanistically determines the upper limit of oscillatory precision. Additionally, these establish ATP not only as a metabolic fuel but also as an effective regulator for cell cycle dynamics, including speed, longevity, and bifurcation.

## ATP depletion reduces both speed and precision of the mitotic oscillator

To investigate cell-cycle performance below endogenous ATP levels, we treated extracts with apyrase, an ATP diphosphohydrolase known to sequentially hydrolyzes ATP to ADP and AMP, and compatible with *Xenopus* cell-free systems^41^. Extreme ATP depletion arrested cell cycle at low Cdk1 states (Fig. 4**a**, top left), while excess ATP drove mitotic arrest with stable high Cdk1 (Fig. 4**a**, top right). These opposing responses highlight the bistable nature of Cdk1 activity and the need to maintain free-energy levels within oscillator network-determined thresholds.

**Fig. 4.**
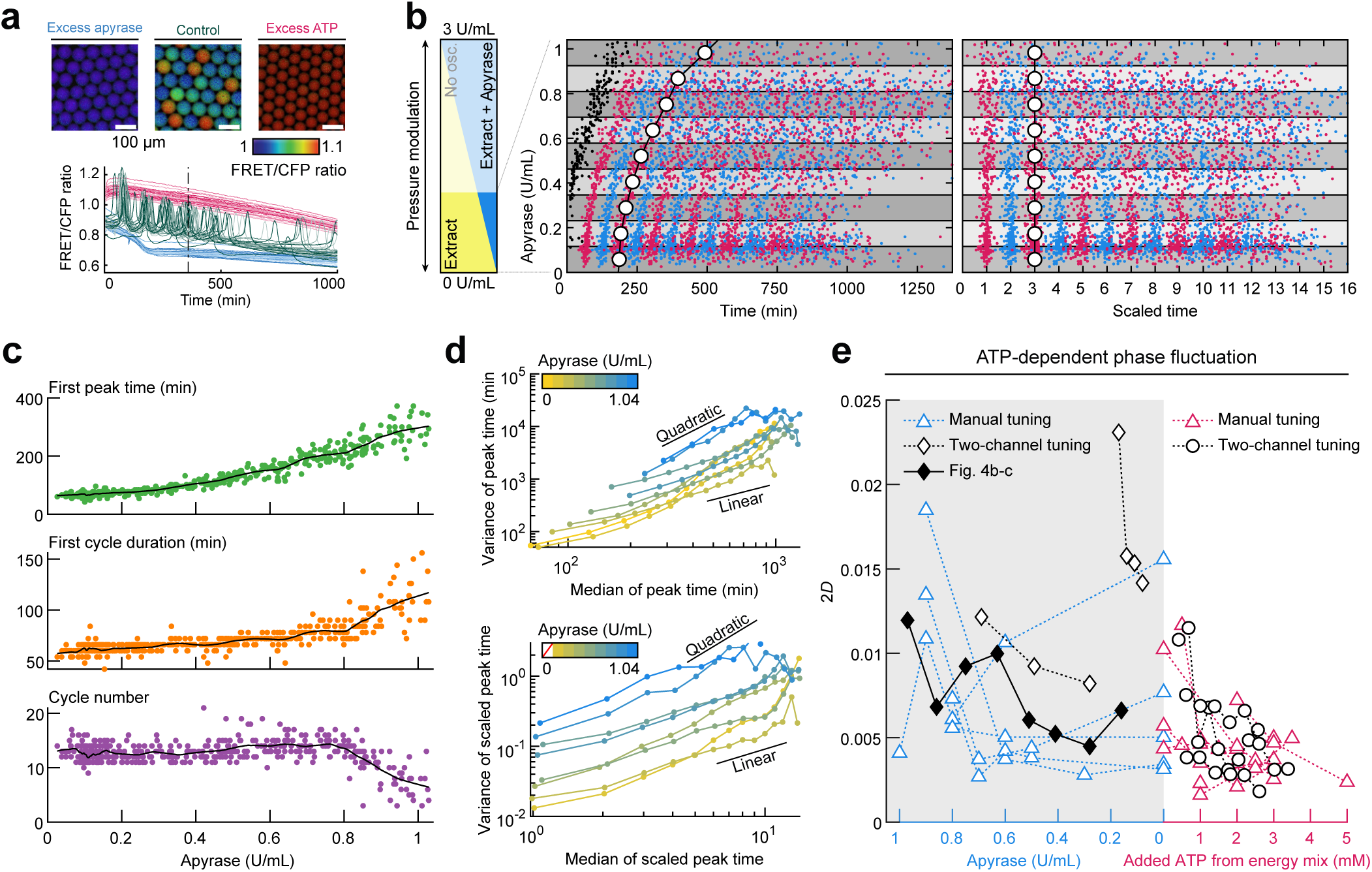
Apyrase-mediated depletion of endogenous ATP reduces the speed and precision of the mitotic oscillator. **a** Cdk1 probe activity indicates distinct cell cycle states under excess apyrase or ATP in the cell-free system. Manually tuned conditions from a single biological replicate are shown with FRET ratio tracks (bottom). The images are colored by the FRET ratio (top) for droplets in each condition at a given time point (black dashed line). Under excessive apyrase (blue) or ATP (magenta) conditions, oscillatory behavior ceased, with Cdk1 arresting at a low or high state, respectively. **b** Two-channel tuning with apyrase. Raster plot of peak times plotted with apyrase levels (left) is shown. The zeroth cycle (black) is ignored for subsequent analysis. Tuning was performed on a range between 0–3 U/mL with no cycles observed for droplets with an estimated apyrase passed 1.2 U/mL. The data was binned into equidistant concentration intervals for subsequent time-rescaling analysis. Time rescaled raster plot is then calculated by interpolation (right). **c** First peak time (top) and first period length (middle) indicate initial offset of the first cycle due to apyrase treatment. Cycle number (bottom) for the data in **b**. **d** Peak variance for the actual (top) peak times and time-rescaled (bottom) peak times of the dataset, both indicating the large dispersion and peak variance associated with apyrase treatment. **e** Our final estimate of the diffusion constant 2*D* including two-channel experiments and manual tuning experiments for apyrase (left side, cyan). Threshold of apyrase dosage was variable across biological replicates. Datasets from Figure 3 plotted for reference (right side, magenta). Stationary criteria of *R*^2^ *>* 0.95 were applied to all datasets.

We created droplets across a broad range of apyrase concentrations (0–3 U/mL; *N* = 443; Fig. 4**b**, left; Supplementary Video 3). QUEEN-2m measurements confirmed apyrase concentration (quantified by dye intensity) correlated with ATP depletion: ratiometric QUEEN-2m signal decreased monotonically with apyrase concentration (Fig. S5**b**), opposite to the trend with ATP addition (Fig. S5**a**). The system was resilient to low depletion (apyrase below 0.4 U/mL), but showed significant period elongation as approaching 1 U/mL, beyond which Cdk1 could not be activated.

To ensure consistent cycle indexing across the ensemble, we excluded the “zeroth” cycle (Fig. 4**b**, left, black; Fig. **S9**), corresponding to the first biological cycle following release from cytostatic factor (CSF) arrest, from our analysis. This cycle was not fully captured in low-apyrase conditions due to the unavoidable time delay required for sample preparation before imaging. This truncation does not affect subsequent *D* estimation, provided that the data represent realizations of a stationary stochastic process. After rescaling and linearity filtering, the processed dataset (Fig. 4**b**, right) revealed progressively delayed first peak time, elongated initial period, and a sharp drop in cycle number at high apyrase concentration (Fig. 4**c**), with sustained oscillations becoming rare above 1 U/mL.

Peak-time variance increased as ATP diminished, evident even from the earliest cycles (Fig. 4**c**, top), and displayed nonlinear growth with cycle index across apyrase conditions (Fig. 4**d**), with higher variance at higher apyrase, consistent with increased heterogeneity across droplets. Using our estimator to offset heterogeneity-induced nonlinear variance, we confirmed that intrinsic oscillatory noise increased monotonically with apyrase concentration (Fig. 4**e**).

Both energy supplementation (Fig. 3) and depletion (Fig. 4) can cause period elongation and bifurcation into the respective high and low stable states of Cdk1 (Fig. 4**a**), with the fastest oscillations occurring at intermediate (near-endogenous) ATP. Their effects on precision were consistent, providing a full dynamic range of energy-precision monotonic dependency. This indicates that the observed noise suppression at higher ATP is not an artifact of rescaling or altered oscillation mechanics, but rather an inherent thermodynamic coupling between dissipation and precision. Dose limits for sustained oscillations varied between apyrase experiments. At the high-apyrase end, oscillations became sparse and showed irregular, pronounced period elongation, likely reflecting a transition toward a critical point, where fluctuations are no longer suppressed by dissipative flux. These underscored the sensitivity of mitotic timing to energetic constraints.

Collectively, these results demonstrate a highly dynamic response of the mitotic oscillator to energy, with both speed and intrinsic noise tightly regulated by cellular free-energy landscapes: while the cell cycle “speed” is non-monotonically tuned by energy, the “precision” is strictly governed by the available energy budget, suggesting that dissipation sets a fundamental limit on the coherence of biological timekeeping.

## Discussion

The early embryonic mitotic oscillator, cleaving a single cell into hundreds with high precision and synchrony, and operating without net biomass accumulation or checkpoint controls, provides an advantageous system for studying the energetics of biological timekeeping. Unlike *in vivo* studies, reconstituted cytoplasmic extracts further isolate the oscillator from complex energy-intensive processes such as DNA replication, nuclear events, actin-dependent mechanical dynamics, chromosome segregation, and cytokinesis, and consequently offer a “minimalist” energetic proxy for the fundamental thermodynamic costs of sustaining Cdk1 oscillations. Our calorimetric measurements (Fig. S10) confirmed that cytoplasmic heat dissipation (∼ 50 nW µL^-1^) represents only a fraction of total embryonic heat production (∼ 0.5 µW µL^-1^)^24,25^, though these measurements still include metabolic processes unrelated to Cdk1 oscillation, and a definitive thermodynamic efficiency bound of the oscillator will ultimately require a fully purified reconstitution of the core oscillator network (Fig. 1**a**).

To quantify cell-cycle coherence with high statistical power, we generated large ensembles of Cdk1 oscillators using droplet microfluidics. A central challenge in quantifying the thermodynamic cost is the quenched randomness from stochastic partitioning during encapsulation^39^, which manifests as period heterogeneity across droplets. We therefore developed an analytical framework to decouple this extrinsic period heterogeneity from intrinsic phase noise, enabling estimation of the diffusion constant *D*^26^. As natural systems encounter similar extrinsic variability, whether from partition errors during cell division or stochastic gene expression, this generalized estimator for *D* is broadly applicable to any heterogeneous oscillator populations across diverse biological systems.

Under endogenous ATP conditions (∼ 3 mM), we estimated *D* on the order of 10^−3^–10^−2^ periods. This estimate has direct implications for the timing of the MBT, a transition from rapid, synchronous cleavages to elongated, desynchronized cycles, regulated by multiple factors, including nuclear-to-cytoplasmic ratio, DNA content, and Chk1 checkpoint signaling^42^. In *Xenopus*, with 12 cleavages before MBT, intrinsic stochasticity alone would accumulate a phase dispersion by 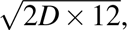 corresponding to 15–49% (4–15 min) of a single 30 min period. Thus, even without active error-correction mechanisms^43^, the basal energy dissipation of the clock appears sufficient to maintain pre-MBT synchrony. This borderline sufficiency implies that the embryonic clock may operate near an energetic optimum, providing adequate precision just enough to prevent premature decoherence while minimizing dissipative cost. This suggests an evolutionary pressure toward metabolic parsimony that prevents the cost of periodicity from being unnecessarily high.

To directly test the energy-precision relationship, we applied multi-channel microfluidics to map the full energy landscape across bifurcation boundaries. In agreement with theory^26^, our results provide direct experimental evidence for an energy-precision trade-off in the mitotic oscillator: increasing the ATP chemical potential monotonically enhances timing precision. The enhanced precision, however, came with a dynamical cost: cell cycle exhibited an intriguing non-monotonic speed dependence on ATP, achieving its fastest oscillation (close to the natural *in vivo* rate) near endogenous concentrations, whereas both ATP depletion and excess caused slowing toward arrest. These observations suggest that the physiological ATP concentration represents an energetic compromise, optimized for maximal oscillation speed while maintaining a baseline precision sufficient for early development.

Although our analysis focuses on stationary limit-cycle behavior, the cell cycle speed is inherently tunable, experiencing programmed slowing during later embryo development and spanning 1.5 orders of magnitude in the *in vitro* Cdk1 oscillator^31,36^. This high degree of period tunability, likely driven by strong positive feedback^44^, is also a natural property of many biological clocks, including heartbeat and neural spikes, which contrasts with the robust near-24-hour period constancy of circadian rhythms. Nonetheless, the qualitative relationship between ATP and precision, mirroring studies in the bacterial Kai circadian clock^26,45–47^, may represent a shared thermodynamic principle across diverse biochemical oscillators. Still, how developmental changes in metabolism and cellular state reshape oscillator precision remains an open question, particularly as differentiating cells encounter heterogeneous energetic environments^48^. Our artificial cell platform, integrating high-throughput droplet generation, precise energetic tuning, and a rigorous, generalizable analysis framework, provides a versatile template for extensive exploration of the stochastic physics in complex biochemical oscillators. These experiments will be useful to understand how biological clocks balance the fundamental trade-offs between speed, precision, and metabolic economy, and test emerging theoretical predictions.

## Methods

### Preparation of *Xenopus* egg extracts

All animal procedures were reviewed and approved by the Institutional Animal Care and Use Committee (IACUC) of the University of Michigan–Ann Arbor (#PRO00011571). Preparation of cell-free cycling extract followed a previously outlined protocol^49^. Sexually competent female *Xenopus laevis* individuals (Nasco and Xenopus1) were primed 3–7 days prior to the experiment with 100 IU of human chorionic gonadotropin (hCG) (MP Biomedicals, 02198591). Frogs were then induced with 500 IU of hCG approximately 12–15 hours before egg collection. Frogs were squeezed for fresh eggs in a 1× Marc’s Modified Ringer (MMR) solution: 5 mM HEPES (Millipore-Sigma, H4034), 100 µM EDTA (Fisher Scientific, BP118), 2 mM CaCl_2_ (MilliporeSigma, C1016), 2 mM KCl (MilliporeSigma, P341), 1 mM MgCl_2_ (MilliporeSigma, M2670), and 100 mM NaCl (MilliporeSigma, 746398), adjusted to pH 7.8 with 10 M NaOH (Fisher Scientific, BP359). The fresh eggs first washed 2-3 times in a 0.2× MMR solution, then they were dejellied with cysteine buffer: 180 mM cysteine (RPI, C50010), 100 mM KCl, 1 mM MgCl_2_, 0.1 mM CaCl_2_, adjusted to pH 7.8 with 10 M NaOH. Dejellied eggs were then treated for activation with 0.5 µg/mL calcium ionophore A23187 (MilliporeSigma, C7522) in 0.2× MMR to release cytostatic factor from meiosis II. The eggs were tightly packed into centrifuge tubes and spun on a swing bucket rotor (Beckman Avanti J-E Centrifuge, JS-13.1 rotor) at 20, 000 × *g* for 10 min. Cytoplasmic section was collected and centrifuged again at 20, 000 × *g* for 5 min for a clean extract product. Cytochalasin B (Cayman Chemical, 11328), leupeptin (Cayman Chemical, 14026), pepstatin A (APExBIO, A2571), chymostatin (Cayman Chemical, 15114) were added at a final concentration of 10 µg/mL each. Interphase-arrested extracts were made with the addition of cycloheximide (MilliporeSigma, C7698) to cycling extracts at a final concentration of 100 µg/mL. The final experimental conditions were made by a 70% extract solution across all experiments including calorimetry, with relevant experimental conditions diluted in 1x extract buffer (EB): 50 mM sucrose (MilliporeSigma, S0389), 10 mM HEPES, 100 mM KCl, 1 mM MgCl_2_, 0.1 mM CaCl_2_, adjusted to pH 7.8 with 10 M NaOH. Buffer solutions for 20× MMR and 20× EB (without sucrose and HEPES) were prepared in bulk and filtered (or autoclaved) for sterility. Final MMR, EB, and cysteine solutions were made fresh on the day of the experiment.

### Preparation of energy mix and apyrase

For a 100× solution of energy mix, we mixed 100 mM ATP (MilliporeSigma, A26209), 10 mM EGTA (MilliporeSigma, E2889), 100 mM MgCl_2_, with experimental trials including/excluding 750 mM creatine phosphate (Chem-Impex, 00072). The pH of the stock was adjusted to between 7.4–7.8 using an Orion^TM^ 9863BN Micro pH and aliquoted to be stored at −80 °C. Apyrase (MilliporeSigma, A6410 or New England Biolabs, M03985) was aliquoted at an estimated 50 U/mL in water and stored at −80 °C.

### Fluorescence-labeled reporters

For measuring the oscillation dynamics of Cdk1, we used the Cdk1-EV-NLS FRET biosensor developed in the lab^31^. Inert recombinant fluorescent proteins (mCherry and BFP) were used as dyes to track concentrations of tuning agents in two-channel experiments. The spectrum of ATP levels generated by the two-channel tuning experiment was validated by QUEEN-2m^40^. All recombinant proteins were over-expressed in BL21 (DE3) bacteria strain under 400 mM IPTG treatment with 20-hour incubation at 18 °C. After harvest, pellets were lysed with lysis buffer containing B-PER (Thermo Scientific, 89821), 1.0 mg/mL lysozyme (MilliporeSigma, L6876), Halt protease inhibitor cocktail (Thermo Scientific, 78430), and Benzonase nuclease (Millipore, E1014) on a rocking shaker for 15 min at room temperature. Cell-lysed buffer was sonicated for complete cell wall disruption. To separate the water-soluble protein and the inclusion body, the samples were centrifuged at 14, 000 × *g* for 15 min at 4 °C. The supernatant was transferred to another tube. All target proteins were tagged with 6× histidine, therefore the supernatant was mixed with the Ni-based resin (Takara Bio, 635660) or His SpinTrap column system (Cytiva, 28401353). Beads were equilibrated with equilibration/wash buffer beforehand. After binding of a target protein and affinity beads, the column was washed three times with the wash buffer. Target proteins were eluted with an elution buffer (300 mM imidazole). The recombinant proteins were dialyzed with Slide-A-Lyzer dialysis cassettes (10,000 MWCO) in PBS overnight and concentrated with Amicon Ultra centrifugal filters. After quantification of protein concentration by the Bradford method, the proteins were diluted to make them 100 µM as a stock concentration. All samples were stored in −80 °C freezer for long-term storage.

### Luciferase assays

Time course luciferase assays were performed on 50 µL bulk extract reactions of 70% cytoplasm solutions and 30% of extract buffer at room temperature. Luciferase reagents CellTiter-Glo 2.0 were used to estimate ATP levels in the extract solution (Promega, G9241). A standard curve was prepared with ATP water solutions prepared between 0.1 µM to 10 µM. ATP solutions were placed on a luminescence well plate with reagents in a 1:1 volume, and 10 min of orbital mixing was set before data acquisition on a Molecular Devices SpectraMax iD3. Extract samples were serially diluted in water to 1:5000 of the original volume and measured on the well plate with a 1:1 ratio of reagent in triplicate as technical replicates of equal volume to the standard curve measurements. SpectraMax was preset with 10 min of orbital mixing before data acquisition for samples and standard curves. After quantifying ATP levels in diluted samples, the endogenous ATP concentration was estimated by dividing by the dilution factor.

### Microfluidics

Microfluidic devices were manufactured using previously described protocols^50^ for SU-8 photolithography on silicon wafers to provide molds for curing of PDMS. The inlets and droplet collection apertures were cut from the cured PDMS slabs peeled off from the molds. A PDMS device was attached to a glass slide coated in a thin layer of partly cured PDMS to fully encapsulate the relevant fluidic circuitry. Pressure modulation of inlets was performed using an Elveflow OB1 MK3+ pressure modulator to generate droplets. A single inlet was used for the surfactant 2% 008-FluoroSurfactant in HFE7500 (Ran Biotechnologies). The surfactant channel splits and recombines with the extract inlet channel to pinch the extract and generate a droplet. Inlet pressure was typically around 2 psi for any particular device. For two-channel experiments we applied a triangular waveform of variable pressure between 0.5 psi to 2.5 psi in the extract inlets to avoid backwash in the microfluidic channels. Some variability in ranges was expected depending on the quality of the device. Hydrophobic-coated glass tubes prepared as described below were used to accommodate a single layer of droplets for imaging. The tubes were placed in a glass-bottom dish and covered in heavy mineral oil for long-term imaging. The image processing of the timelapse data was as previously described^50^.

### Two-channel and manual tuning

Two distinct designs of microfluidic devices were used. Single-channel devices were used for generating droplets from a single homogeneous extract solution (Fig. 1**a**). Manual tuning was achieved by preparing separate homogeneous solutions for specific experimental conditions of interest. Alternatively, a two-channel device with two extract inlets (Fig. 3**a**) was used to collect data over a wide range of concentrations for a single tuning parameter. In these experiments, two separate homogeneous solutions were mixed in the microfluidic device by varying the extract inlet pressure in an oscillatory manner that was out of phase. We thereby performed a high-throughput generation of extract droplets with intermediate concentrations of the tuning parameter. One of the tuning solutions included a fluorescent dye to track the tuning parameter in the mixed droplet population. To map concentration to fluorophore intensity in a two-channel tuning setup, fixed-pressure droplet generation was performed at each of the two homogeneous concentrations.

### Glass tube coating

As previously described^32^, we used a VitroCom miniature hollow glass tubing with a wall thickness of 100 µm (VitroCom, 5012). A heating block was heated to 95 °C in an incubator and then placed in a polycarbonate vacuum desiccator with a white polypropylene bottom (Bel-Art, F42025-0000). Glass tubes and a 1.5 ml Eppendorf tube containing 30 µL Trichloro(1H,1H,2H,2H-perfluorooctyl)silane (MilliporeSigma, 448931) were placed in the heating block. Vacuum was applied to the desiccator for approximately 1 min. Then the valve was closed, and the tubes were left incubated overnight. The coated glass was then cut into pieces with lengths of 3–5 mm, bathed in 70% ethanol, and left to dry before experimental use.

### Time-lapse fluorescence microscopy

Time-lapse imaging was performed with an inverted microscope IX83 (Olympus) equipped with a UPlanSApo 4x/0.16 objective lens (Olympus), an ORCA-Flash4.0 V3 digital CMOS camera (Hamamatsu Photonics), X-Cite Xylis broad spectrum LED illumination system (Exelitas Technologies), and a motorized XY stage (Prior Scientific) at room temperature. The microscope was controlled by Micro-Manager software^51^. Time-lapse images were recorded in bright-field and multiple fluorescence channels for up to 2 days at a time interval of 3–6 min, and dynamics of individual droplets were analyzed using a custom semi-automated segmentation and tracking algorithm^36,50^,

### Image analysis and processing

Images were analyzed using custom MATLAB (MathWorks) scripts developed in a previous study with some novel optimizations^50^. Bright field images were segmented using the watershed algorithm to identify droplets. Threshold criteria for eccentricity and size were applied to each segment to isolate droplet data, and segments were tracked through the course of the experiment. Segments were used to extract fluorescence intensity for all imaged channels. FRET/CFP emission ratio for Cdk1 activity was calculated by dividing the mean of FRET channel intensity by the mean of CFP channel intensity for individual droplets. A custom peak selection algorithm was applied to the FRET/CFP ratio to identify Cdk1 oscillations. After automated peak selection and a visual inspection/correction of all peak detection, we redefine which of the detected peaks are considered the first peak of the experiment (Fig. S9). The resulting datasets were then used for precision analysis.

### Precision quantification

Further information is available in Supplementary Information.

### Real-Time quantitative PCR

The total RNA was extracted from the bulk extract using the TRIzol reagent (Invitrogen) and the isopropanol precipitation method with GlycoBlue (Invitrogen). In summary, 1 µL of the bulk extract was dissolved in 1 mL of TRIzol reagent, then total RNA was extracted according to the product user guide. TRIzol containing extract was mixed with chloroform, vigorously vortexed for 15 s, and incubated for 2 min at room temperature. The aqueous phase was separated by centrifugation at 12, 000 × *g* for 15 min at 4 °C, then transferred to a new tube containing 600 µL ice-cold isopropanol and 1.5 µL of GlycoBlue co-precipitant, and mixed well by inverting the tube. After 10 min room temperature incubation, the blue pellet was formed by centrifugation at 12, 000 × *g* for 10 min at 4 °C. The supernatant was discarded as much as possible without disturbing the RNA-containing blue pellet. The pellet was washed with 75% ethanol and centrifuged at 7, 500 × *g* for 10 min at 4 °C. After discarding the supernatant, the sample was completely air-dried in the hood. The total RNA was solubilized with 30 µL of RNase-free water, and its concentration and purity were measured by a spectrophotometer. cDNA was synthesized with iScript (Bio-Rad) using the manufacturer’s instructions. 500 ng of total DNA was used for cDNA synthesis in a total 20 µL reaction. The random primers in the reagent ensure that the first strand synthesis includes total RNA, including mRNA and rRNA. The synthesized cDNA sample was transferred to a 1.5 mL tube and stored at -80 °C freezer until use. To generate a standard curve at RT-qPCR, *Xenopus laevis* cyclin B1 S homolog gene **ccnb1.S** (NCBI Genbank ID: NM 001088520.1) sequence was synthesized (Twist Bioscience) and cloned into the pCS2(+) vector in the lab. Its mRNA was transcribed by mMESSAGE mMACHINE SP6 mRNA kit (Invitrogen) and purified by RNeasy micro kit (Qiagen). RT-qPCR primers were designed using the Custom TaqMan Assay Design Tool provided by the ThermoFisher Scientific website. The 150–200 bp of the target mRNA region was selected from the exon-to-exon region, where possible to avoid genomic DNA amplification. The input data is the coding DNA sequence (CDS) of NM 001088520.1. The web tool automatically designed both forward and reverse primers and a probe with FAM dye. Real-time quantitative PCR (RT-qPCR) was performed using the CFX96 system (Bio-Rad) on 96 well plates with a standard reaction per well containing 1/10 diluted cDNA as a template (2 µL per well), TaqMan Gene Expression Master Mix (10 µL per well), Custom TaqMan Assay (1 µL per well), and RNase-free water (7 µL per well). The RT-qPCR reaction for each sample was conducted in duplicates or triplicates (technical replicate). Water and no-template controls were used as negative controls for each primer set. The RT-qPCR data were analyzed using the CFX Manager software. Cycle threshold (Ct) values were obtained using auto baseline and applied to all amplicons of the same primer set. Standard curve and mRNA level estimation were calculated using a custom MATLAB code.

### Cell-free isothermal calorimetry

The heat dissipation estimates for cytoplasmic *Xenopus laevis* cycling extract were extrapolated from isothermal calorimetry data. Solutions of bulk cytoplasmic extract were prepared to match the microscopy conditions. A dilution mixture of 1× extract buffer and the respective levels of energy mix was degassed for 10 min, then added to cytoplasm extract for a 30% dilution. A volume of 300 µL cytoplasmic solution was then inserted in a low-volume Nano ITC (TA Instruments) set at 22 °C. Degassed water of equal volume was placed in the reference cell. In stirring experiments, the reference cell plunger and the injection syringe were left on, with the syringe plunger removed. The injection protocol was neglected. In static extract experiments the syringe and the reference cell plunger were removed. The baseline did not auto-equilibrate for cytoplasmic extract. A separate run with water was used in the sample cell for a baseline subtraction.

## Acknowledgments

We thank the current and past members of the Yang lab for the scientific discussions and feedback on the manuscript; Yuansheng Cao and Jordan Horowitz for their input and discussion over the statistical analysis; Hiyoyuki Noji for providing pRSET-QUE2m (Addgene plasmid #129350); Owen Puls, and Minjun Jin for assistance with the preparation of *Xenopus* egg extracts and the droplet-based microfluidic devices. This work was supported by the National Science Foundation (MCB#2218083), the National Institutes of Health (R01GM144584), and the Margaret and Herman Sokol Faculty Awards at the University of Michigan to Q.Y., and an NSF Graduate Research Fellowship (GRFP) to L.Y.

## Author Contributions

QY and SW conceived the project. QY supervised the project. SW, LY, and GM performed experiments, image analysis, and analyzed data. YK developed drift correction analysis. SW, LY, GM, YK, DW, QY helped with data interpretation. LY, YK, GM, QY wrote the manuscript. SW, LY, GM, YK, UK, QY, edited the manuscript.

## Data Availability

There is no restriction on data availability. All datasets necessary to reproduce the figures in the manuscript will be deposited in a public repository upon publication. Source data will be provided with the paper. Additional raw imaging data are available from the corresponding author upon reasonable request.

## Code Availability

There is no restriction on code availability. MATLAB scripts for droplet analysis are deposited on GitHub.

## References

1. Marrable, A. Variations in early cleavage of the zebra fish. Nature 184, 1160–1161 (1959).

2. Olivier, N. et al. Cell lineage reconstruction of early zebrafish embryos using label-free nonlinear microscopy. Science 329, 967–971 (2010).

3. Tsai, T. Y.-C., Theriot, J. A. & Ferrell Jr, J. E. Changes in oscillatory dynamics in the cell cycle of early Xenopus laevis embryos. PLoS Biol. 12, e1001788 (2014).

4. Kim, S. Y., Song, E. J., Lee, K.-J. & Ferrell Jr, J. E. Multisite M-phase phosphorylation of Xenopus Wee1A. Mol. Cell. Biol. 25, 10580–10590 (2005).

5. Trunnell, N. B., Poon, A. C., Kim, S. Y. & Ferrell, J. E. Ultrasensitivity in the regulation of Cdc25C by Cdk1. Mol. Cell. 41, 263–274 (2011).

6. Pomerening, J. R., Sontag, E. D. & Ferrell Jr, J. E. Building a cell cycle oscillator: hysteresis and bistability in the activation of Cdc2. Nat. Cell. Biol. 5, 346–351 (2003).

7. Ferrell, J. E., Tsai, T. Y.-C. & Yang, Q. Modeling the cell cycle: why do certain circuits oscillate? Cell 144, 874–885 (2011).

8. Yang, Q. & Ferrell Jr, J. E. The Cdk1–APC/C cell cycle oscillator circuit functions as a time-delayed, ultrasensitive switch. Nat. Cell. Biol. 15, 519–525 (2013).

9. Guan, Y., Wang, S., Jin, M., Xu, H. & Yang, Q. Reconstitution of cell-cycle oscillations in microemulsions of cell-free Xenopus egg extracts. J. Vis. Exp. 58240 (2018).

10. Li, D., Liao, S., Ouyang, Q. & Li, F. Interplay between atp and hydrolysis free energy in promoting and restraining biochemical switches. Phys. Rev. Res. 6, 033050 (2024).

11. Miniowitz-Shemtov, S., Teichner, A., Sitry-Shevah, D. & Hershko, A. ATP is required for the release of the anaphase-promoting complex/cyclosome from inhibition by the mitotic checkpoint. Proc. Natl. Acad. Sci. 107, 5351–5356 (2010).

12. Peth, A., Uchiki, T. & Goldberg, A. L. ATP-dependent steps in the binding of ubiquitin conjugates to the 26S proteasome that commit to degradation. Mol. Cell. 40, 671–681 (2010).

13. Peth, A., Nathan, J. A. & Goldberg, A. L. The ATP costs and time required to degrade ubiquitinated proteins by the 26 S proteasome. J. Biol. Chem. 288, 29215–29222 (2013).

14. Holt, L. J. et al. Global analysis of Cdk1 substrate phosphorylation sites provides insights into evolution. Science 325, 1682–1686 (2009).

15. Petrone, A., Adamo, M. E., Cheng, C. & Kettenbach, A. N. Identification of candidate cyclin-dependent kinase 1 (Cdk1) substrates in mitosis by quantitative phosphoproteomics. Mol. & Cell. Proteomics 15, 2448–2461 (2016).

16. Massacci, G., Perfetto, L. & Sacco, F. The cyclin-dependent kinase 1: more than a cell cycle regulator. Br. J. Cancer 129, 1707–1716 (2023).

17. Lane, N. & Martin, W. The energetics of genome complexity. Nature 467, 929–934 (2010).

18. Kimura, K. & Hirano, T. ATP-dependent positive supercoiling of DNA by 13S condensin: a biochemical implication for chromosome condensation. Cell 90, 625–634 (1997).

19. Collas, P. Sequential PKC-and Cdc2-mediated phosphorylation events elicit zebrafish nuclear envelope disassembly. J. Cell. Sci. 112, 977–987 (1999).

20. Hetzer, M. et al. Distinct AAA-ATPase p97 complexes function in discrete steps of nuclear assembly. Nat. Cell. Biol. 3, 1086–1091 (2001).

21. Zhang, X., Wu, X. Q., Lu, S., Guo, Y. L. & Ma, X. Deficit of mitochondria-derived atp during oxidative stress impairs mouse mii oocyte spindles. Cell. Res. 16, 841–850 (2006).

22. Ott, C. et al. VPS4 is a dynamic component of the centrosome that regulates centrosome localization of γ-tubulin, centriolar satellite stability and ciliogenesis. Sci. Rep. 8, 3353 (2018).

23. Arumugam, P. et al. ATP hydrolysis is required for cohesin’s association with chromosomes. Curr. Biol. 13, 1941–1953 (2003).

24. Nagano, Y. & Ode, K. L. Temperature-independent energy expenditure in early development of the african clawed frog Xenopus laevis. Phys. Biol. 11, 046008 (2014).

25. Rodenfels, J., Neugebauer, K. M. & Howard, J. Heat oscillations driven by the embryonic cell cycle reveal the energetic costs of signaling. Dev. Cell. 48, 646–658 (2019).

26. Cao, Y., Wang, H., Ouyang, Q. & Tu, Y. The free-energy cost of accurate biochemical oscillations. Nat. Phys. 11, 772–778 (2015).

27. Barato, A. C. & Seifert, U. Thermodynamic uncertainty relation for biomolecular processes. Phys. Rev. Lett. 114, 158101 (2015).

28. Zotin, A., Faustov, V., Radzinskaja, L. & Ozernyuk, N. ATP level and respiration of embryos. Development 18, 1–12 (1967).

29. Morgan, D. O. The Cell Cycle: Principles of Control. Primers in Biology (New Science Press Ltd, 2007).

30. Murray, A. W. Cell cycle extracts. Methods Cell. Biol. 36, 581–605 (1991).

31. Maryu, G. & Yang, Q. Nuclear-cytoplasmic compartmentalization of cyclin B1-Cdk1 promotes robust timing of mitotic events. Cell. Rep. 41 (2022).

32. Guan, Y. et al. A robust and tunable mitotic oscillator in artificial cells. eLife 7, e33549 (2018).

33. Miller, D. & Horowitz, S. Intracellular compartmentalization of adenosine triphosphate. J. Biol. Chem. 261, 13911–13915 (1986).

34. Woodland, H. & Pestell, R. Determination of the nucleoside triphosphate contents of eggs and oocytes of Xenopus laevis. Biochem. J. 127, 597–605 (1972).

35. Valentine, M., Perlman, Z., Mitchison, T. & Weitz, D. Mechanical properties of xenopus egg cytoplasmic extracts. Biophys. J. 88, 680–689 (2005).

36. Li, Z. et al. Comprehensive parameter space mapping of cell cycle dynamics under network perturbations. ACS Synth. Biol. 13, 804–815 (2024).

37. Gillespie, D. T. A general method for numerically simulating the stochastic time evolution of coupled chemical reactions. J. Comput. Phys. 22, 403–434 (1976).

38. Huh, D. & Paulsson, J. Random partitioning of molecules at cell division. Proc. Natl. Acad. Sci. 108, 15004–15009 (2011).

39. Weitz, M. et al. Diversity in the dynamical behaviour of a compartmentalized programmable biochemical oscillator. Nat. Chem. 6, 295–302 (2014).

40. Yaginuma, H. et al. Diversity in atp concentrations in a single bacterial cell population revealed by quantitative single-cell imaging. Sci. Rep. 4, 6522 (2014).

41. Buendia, B., Draetta, G. & Karsenti, E. Regulation of the microtubule nucleating activity of centrosomes in Xenopus egg extracts: role of cyclin A-associated protein kinase. The J. Cell. Biol. 116, 1431–1442 (1992).

42. Farrell, J. A. & O’Farrell, P. H. From egg to gastrula: how the cell cycle is remodeled during the Drosophila mid-blastula transition. Annu. Rev. Genet. 48, 269–294 (2014).

43. Anderson, G. A., Gelens, L., Baker, J. C. & Ferrell, J. E. Desynchronizing embryonic cell division waves reveals the robustness of Xenopus laevis development. Cell. Rep. 21, 37–46 (2017).

44. Tsai, T. Y.-C. et al. Robust, tunable biological oscillations from interlinked positive and negative feedback loops. Science 321, 126–129 (2008).

45. Barkai, N. & Leibler, S. Circadian clocks limited by noise. Nature 403, 267–268 (2000).

46. Rust, M. J., Golden, S. S. & O’Shea, E. K. Light-driven changes in energy metabolism directly entrain the cyanobacterial circadian oscillator. Science 331, 220–223 (2011).

47. Phong, C., Markson, J. S., Wilhoite, C. M. & Rust, M. J. Robust and tunable circadian rhythms from differentially sensitive catalytic domains. Proc. Natl. Acad. Sci. 110, 1124–1129 (2013).

48. Greiner, J. V. & Glonek, T. Intracellular atp concentration and implication for cellular evolution. Biology 10, 1166 (2021).

49. Jin, M., Tavella, F., Wang, S. & Yang, Q. In vitro cell cycle oscillations exhibit a robust and hysteretic response to changes in cytoplasmic density. Proc. Natl. Acad. Sci. 119, e2109547119 (2022).

50. Sun, M., Li, Z., Wang, S., Maryu, G. & Yang, Q. Building dynamic cellular machineries in droplet based artificial cells with single-droplet tracking and analysis. Anal. Chem. 91 (2019).

51. Edelstein, A. D. et al. Advanced methods of microscope control using µManager. J. Biol. Methods 1, e10 (2014).

